# Holstein Friesian dairy cattle edited for diluted coat color as adaptation to climate change

**DOI:** 10.1101/2020.09.15.298950

**Authors:** G. Laible, S-A. Cole, B. Brophy, J. Wei, S. Leath, S. Jivanji, M.D. Littlejohn, D.N. Wells

## Abstract

High-producing Holstein Friesian dairy cattle have a characteristic black and white coat pattern where black frequently comprises a large proportion of the coat. Compared to a light coat color, black absorbs more solar radiation translating into radiative heat gain which is a contributing factor to heat stress in cattle, negatively impacting on their production levels, fertility and welfare. To better adapt dairy cattle to the rapidly changing climatic conditions with predictions for more frequent and prolonged hot temperature patterns, we aimed to lighten their coat color by genome editing. Using gRNA/Cas9-mediated editing, we introduced a three base pair (bp) deletion in the pre-melanosomal protein 17 gene (*PMEL*) proposed as the causative variant responsible for the semi-dominant color dilution phenotype seen in Galloway and Highland cattle. Calves generated from cells homozygous for the edited mutation revealed a strong color dilution effect. Instead of the characteristic black and white coat color patterning of control calves generated from unedited parental cells, the edited calves displayed a novel pattern of grey and white markings and absence of any black areas. This, for the first time, verified the causative nature of the *PMEL* mutation for diluting the black coat color in cattle. With these edited animals, it is now possible to dissect the effects of the introgressed edit and other interfering allelic variants that might exist in individual cattle and accurately determine the impact of only the three bp change on important health, welfare and production traits. In addition, our study proved targeted editing as a promising approach for the rapid adaptation of livestock to changing climatic conditions.

## Introduction

The present trend of increasing global temperatures are rapidly changing the environment and conditions under which dairy animals are grazing. Hence, dairy cattle are no longer well adapted to the predicted realities of more frequent and prolonged periods of hotter summer temperatures (1–4). This poses significant challenges for their welfare and negatively impacts on their eco-productivity (5–7). Already, New Zealand dairy cows become heat stressed for close to 20% of lactation days in major dairy regions in New Zealand (8). This is particularly relevant for animals with black hair, a common characteristic of Holstein Friesian dairy cattle, which absorb twice as much solar radiation as white hair (9). Hence, it exposes black animals to enhanced radiative heat gain which contributes to heat stress. In hot weather, this results in a reduced ability of primarily black dairy cows to regulate body temperature and maintain milk production levels (10) while also negatively impacting on their reproductive performance (11). Lightening of the coat color should help to reduce these impacts and provide a first step to better adapted dairy cattle. In different species, *PMEL* mutations were shown to be responsible for color dilution effects (12–17). A deletion of a leucine codon in the signal peptide (p.Leu18del) of PMEL segregates in Highland, Galloway, and Tuli cattle, and has been proposed as a causative mutation for coat color effects (18, 19). Animals heterozygous for the p.Leu18del mutation display a faded version of the wild type (WT) black or red coat color, referred to as dun or yellow, respectively. Homozygotes present an even stronger dilution effect with a white to off-white coat, also called silver dun (black WT) or white (red WT) (18). While the role of this variant has not been functionally confirmed, its proposed coat color impacts make it an excellent candidate for introgression into Holstein Friesian dairy cattle to reduce radiative heat gain and improve overall heat tolerance.

Although the introgression of the p.Leu18del mutation would be possible with a conventional crossbreeding strategy, such an approach would not enable immediate functional validation of the p.Leu18del variant since many other variants on the same chromosome are co-inherited when crossbreeding. In addition, it would require back crossing over many years to catch up to the genetic merit of contemporaneous animals, which renders this process unsuitably slow for a timely adaptation. By contrast, genome editing can prove the causative relationship between a specific mutation and impact on phenotype, and provide the scope for rapid introgression - essentially within a single generation (20, 21).

In this study, we report the gRNA/Cas9-mediated introgression of the naturally occurring mutation p.Leu18del in the *PMEL* gene known from Galloway and Highland cattle, into Holstein Friesian cattle. Calves homozygous for the *PMEL* mutation displayed a distinct color dilution to a silvery grey of the otherwise black coat markings in the control calves. The white areas remained unaffected. This novel grey and white coat color phenotype demonstrates the causative role of the *PMEL* mutation in coat color dilution. In addition, our study shows that the introgression of naturally occurring sequence variants by genome editing is a promising new approach to rapidly improve and adapt livestock to changing environmental conditions.

## Materials and Methods

### Animal studies

All animal experiments were performed in accordance with the relevant guidelines and regulations with approvals from New Zealand’s Environmental Protection Authority (GMD100279) and the Ruakura Animal Ethics Committee (14236).

### Signal peptide prediction

Signal peptide features of WT and deletion variant were evaluated using Signal P-5.0 (22).

### Genome editing

*PMEL*-specific gRNAs were designed using the Crispor online tool (23). The plasmid pX330 was modified by inserting high scoring gRNA sequences (Table 1) as previously described (24, 25). These expression plasmids were subsequently used to simultaneously deliver Cas9 nuclease and gRNA into male primary bovine fetal fibroblast (BEF2) cells. BEF2 cells were routinely cultured in Dulbecco’s Modified Eagle’s Medium (DMEM)/F12 supplemented with 10% fetal calf serum (Moregate Biotech). For editing, 2×10^5^ BEF2 cells were transfected with a gRNA/Cas9 plasmid (0.5 μg) or co-transfected with a gRNA/Cas9 plasmid (0.5 μg) plus homology-directed repair (HDR) template (0.4 μg) using a 10 μl tip with program A3 (1500 V pulse voltage, 20 msec pulse width, 1 pulse) according to the manufacturer’s instruction of the Neon transfection system (Invitrogen). Following transfection, cells were reseeded and cultured for 2 days. After gently tapping the culture plate, dislodged mitotic cells were manually picked with a glass capillary and transferred into individual wells of a 96 well plate for the isolation of cell clones. After approximately 14 days cell clones reached confluency and were successively transferred to larger multi-well plate formats for further expansion and cryopreservation.

**Table 1.**
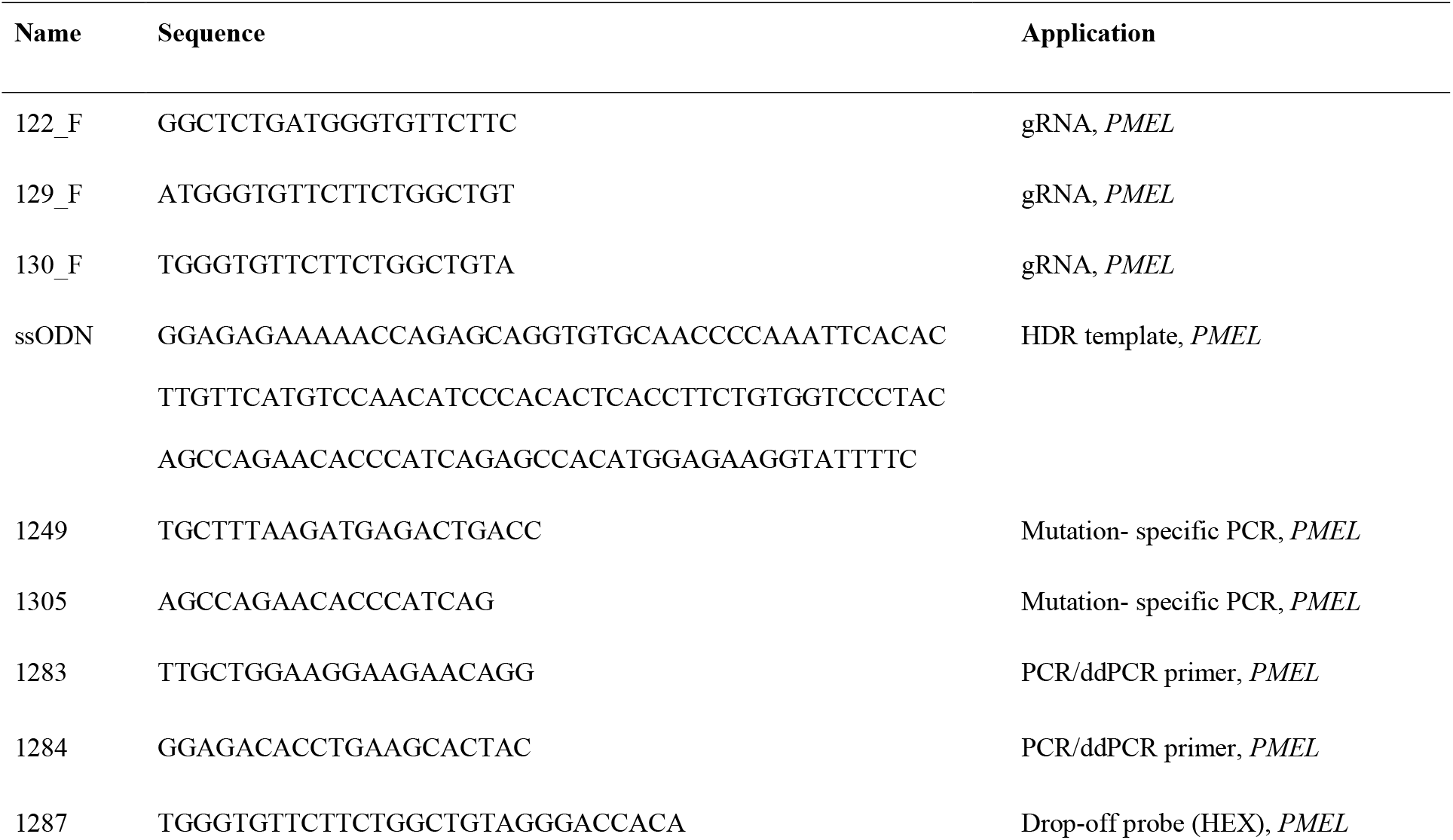
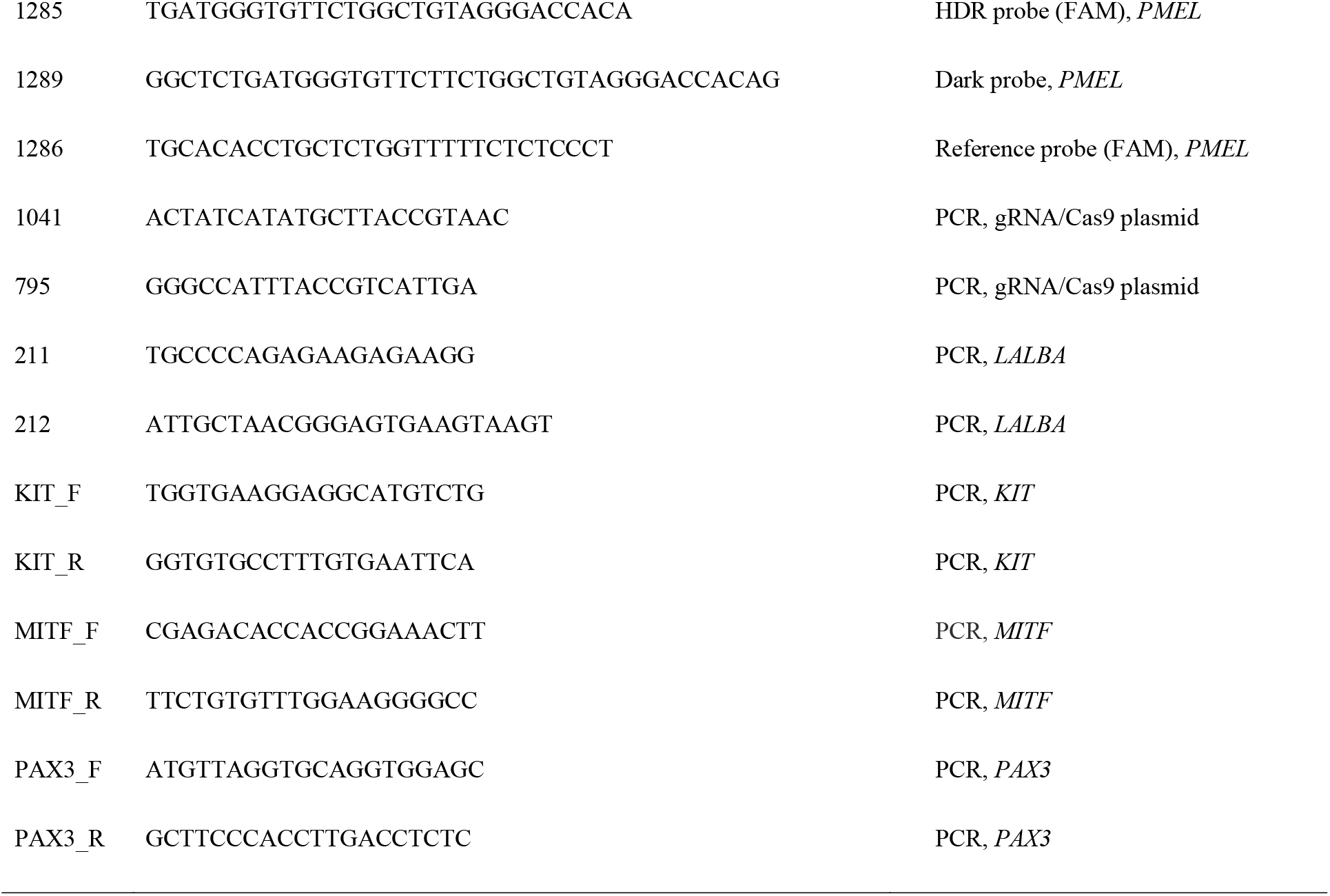
PCR primer, probe, gRNA and repair template sequences used to characterise the *PMEL* locus and white spotting genes.

### Genotype analysis

Genomic DNA was extracted from cultured bovine cells and blood samples as previously described (26). Sequences of all PCR primers used in this study are listed in Table 1. The relevant region of the *PMEL* locus was PCR amplified with primer pair 1283/1284 using a Kapa 2G Fast Hotstart PCR kit (KapaBiosystems). Cycling conditions were 95 °C, 3 min; 95 °C, 15 s, 60 °C, 15 s, and 72 °C, 1 s for 35 cycles; and a final extension step of 72 °C for 5 min. For the mutation-specific PCR, the primer pair was 1249/1305 and used the same amplification conditions as above. Similarly, the same conditions were used for the PCR-genotyping of the three ‘white spotting’ tag variants with primers representing the *KIT* (KIT_F/R), *MITF* (MITF_F/R), and *PAX3* (PAX3_F/R) loci of interest.

Probing for potential gRNA/Cas9 plasmid integration was performed with the vector-specific primers 1041 and 795 amplifying a 489 bp fragment from the U6 promoter across the gRNA and into the CMV enhancer. Amplification of a 444 bp fragment from the endogenous bovine alpha-lactalbumin gene (*LALBA*) with primers 211 and 212 served as a control for the presence of amplifiable DNA. Cycle conditions were as detailed above.

Digital droplet PCR (ddPCR) assays were performed on a QX200 Droplet Digital PCR System (Bio-Rad) according to the method detailed in the Bio-Rad Droplet Digital™ PCR Applications Guide essentially as described before (27). Briefly, ddPCR assays for non-homologous end joining (NHEJ) events were performed with approximately 100 ng of genomic DNA as template, primer pair 1283/1284, and a HEX-labelled drop-off (1287) and FAM-labelled reference probe (1286) located on the same amplicon. For determining template-mediated HDR editing, the drop-off probe was replaced by an HDR-probe (1285) and an unlabelled dark probe (1289). Following droplet generation, samples were PCR amplified using a Bio-Rad PCR machine as follows: 95 °C for 10 min, followed by 40 cycles of 94 °C 30 sec, 60 °C 1 min (ramp rate 2 °C/sec), then 98 °C for 10 min. Amplification results were acquired with a Bio-Rad Droplet Reader and analyzed using QuantaSoft™ Analysis Pro Software (Bio-Rad).

For sequence analyses, PCR reactions were separated on agarose gels and PCR fragments isolated using a Nucleo-Spin Gel and PCR Clean-up kit (Macherey-Nagel). Sanger sequencing of *PMEL* and *KIT, MITF* and *PAX3* fragments was provided by Massey Genome Service (Palmerston North, New Zealand) and the Auckland Genomics facility at the University of Auckland (Auckland, New Zealand), respectively. Tracking of Indels by DEcomposition (TIDE, Brinkman et al., 2014) was used where required to dissect sequences of different gRNA/Cas9-generated mutations intertwined in the sequencing result of the amplified target region.

### Somatic cell nuclear transfer

SCNT embryos were reconstructed from abattoir-derived, enucleated oocytes and individual, serum starved-donor cells using a zona-free cloning procedure as previously described (26). After seven days in vitro culture single embryos were non-surgically transferred to synchronized recipient cows for development to term.

### Hair and skin color measurements

The lightness of color was measured using a Hunter Miniscan XE colorimeter (HunterLab, Hunter Associates Laboratories Inc., Reston, USA) according to the CIELAB color system (28). The lightness of color is measured as an L* value that is defined by the position on the black (0) to white (100) axis of the visible spectrum. For each of the three control calves, the L* values were determined for the hair of two different white and two dark markings, with one area located on the right and the other on the left flank of each animal. To determine the corresponding values for skin, the hair from a small area in these markings was shaved and L* values were measured for two light and two dark markings. For the PMEL mutant calf, the coat/pelt was removed following the death of the animal and used for the color measurements. Again, two white and two dark markings on opposing flanks were selected and L* determined with three individual measurements each.

### Statistical analysis

Statistical significance levels of observed differences were determined by the two-tailed Fisher exact test for independence in 2 x 2 tables (cloning efficiency of CC14 vs WT) and two-tailed student’s t-test (L* measurements).

## Results

### gRNA/Cas9-mediated editing of the p.Leu18del *PMEL* mutation

The three bp deletion *PMEL* variant, resulting in the deletion of leucine 18 in the signal peptide of PMEL, is known from Highland cattle where it is associated with a semi-dominant color dilution phenotype (18).

To introgess this naturally occurring sequence variant into Holstein Friesian cattle we first designed three different gRNAs targeting the *PMEL* gene sequence near the mutation site. The corresponding gRNA/Cas9 editors were then evaluated by ddPCR for target-specific cleavage activity and efficiency for template-directed repair. The analysis of primary bovine embryonic fibroblast cells (BEF2) individually transfected with the gRNA/Cas9 editors 122_F, 129_F and 130_F showed that all three editors could generate indel mutations at the target site following NHEJ repair (Table 2). The best mutation rate was attained using gRNA/Cas9 editor 129_F with 35 % of sequences containing indel mutations. To measure HDR efficiency we co-transfected BEF2 cells with the gRNA/Cas9 editors and a 127 bp single stranded oligonucleotide specifying the three bp deletion of the p.Leu18del PMEL mutation (Figure 1). This confirmed that the highest HDR activities were associated with gRNA/Cas9 editors 122_F and 129_F and resulted in 5.7 % and 6.0 % of HDR events, respectively in transfected BEF2 cells.

**Figure 1.**
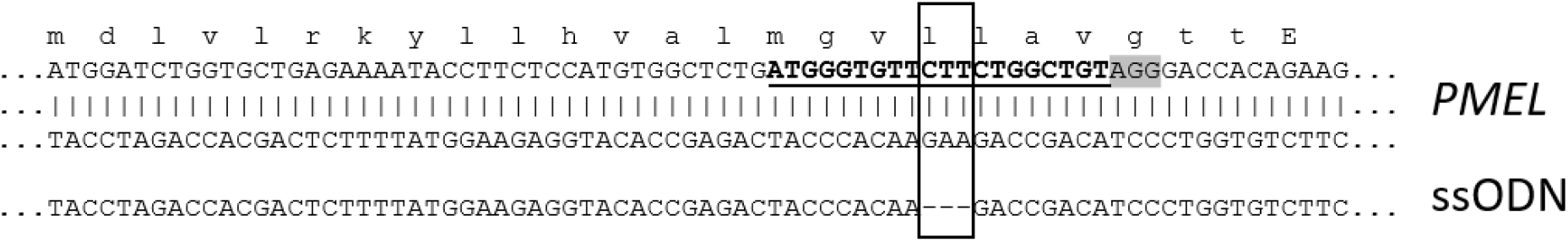
Schematic overview of the *PMEL* target region. Shown is the DNA sequence for the relevant region of the *PMEL* gene with the binding site for editor 129_F (bold, underlined), PAM sequence (grey highlight) and the location of the three bp deletion (box). Below is the aligned sequence of the single stranded HDR template (ssODN) specifying a three bp deletion (dashes). The corresponding amino acid sequence for PMEL is given in single letter code above the DNA sequence with lower case indicating amino acids of the predicted signal peptide according to the UniProt annotation for bovine PMEL (entry Q06154).

**Table 2.** Target-specific editing activity of three PMEL-specific gRNA/Cas9 editors.

| gRNA/Cas9 editor | % NHEJ | % HDR |
| --- | --- | --- |
| 122_F | 11 | 5.7 |
| 129_F | 35 | 6.0 |
| 130_F* | 7 | 3.8 |

\*based on a single transfection

### Isolation of precisely edited cell clones

Based on our initial testing, we selected gRNA 129_F for the gRNA/Cas9-mediated editing of BEF2 cells for the isolation of edited cell clones. Following co-transfection of the PMEL-specific editors and HDR template, we picked 128 individual mitotic cells into 96 well plates for clonal expansion. Of those, 96 expanded into cell clones that were characterised (Table 3). The isolated cell clones were first screened by mutation-specific PCR, that was designed to only amplify the three bp deletion allele. This initial screening was done at low stringency to minimize the risk of failing to detect some correctly edited cell clones. It identified 30 candidate clones for the intended HDR edit. These candidates were then more stringently screened by ddPCR using a hydrolysation probe specific for the three bp HDR mutation. This revealed a total of seven clones that had at least one HDR allele, and in combination with sequencing and TIDE analysis, three clones were confirmed that had one HDR and one WT allele (monoallelic) and four clones that were biallelically edited with two HDR alleles (Figure 2).

**Table 3.** Summary of cell clone isolation.

| Isolation/Characterisation Step | No. of Cell Clones | Efficiency |
| --- | --- | --- |
| Cell clones picked | 128 | 100% |
| Expanded cell clones | 96 | 75% (96/128) |
| Mutation specific PCR +ve | 30 | 31% (30/96) |
| ddPCR HDR +ve, total | 7 | 7% (13/96) |
| ddPCR HDR +ve, monoallelic | 3 | 3% (3/96) |
| ddPCR HDR +ve, biallelic | 4 | 4% (4/96) |

**Figure 2.**
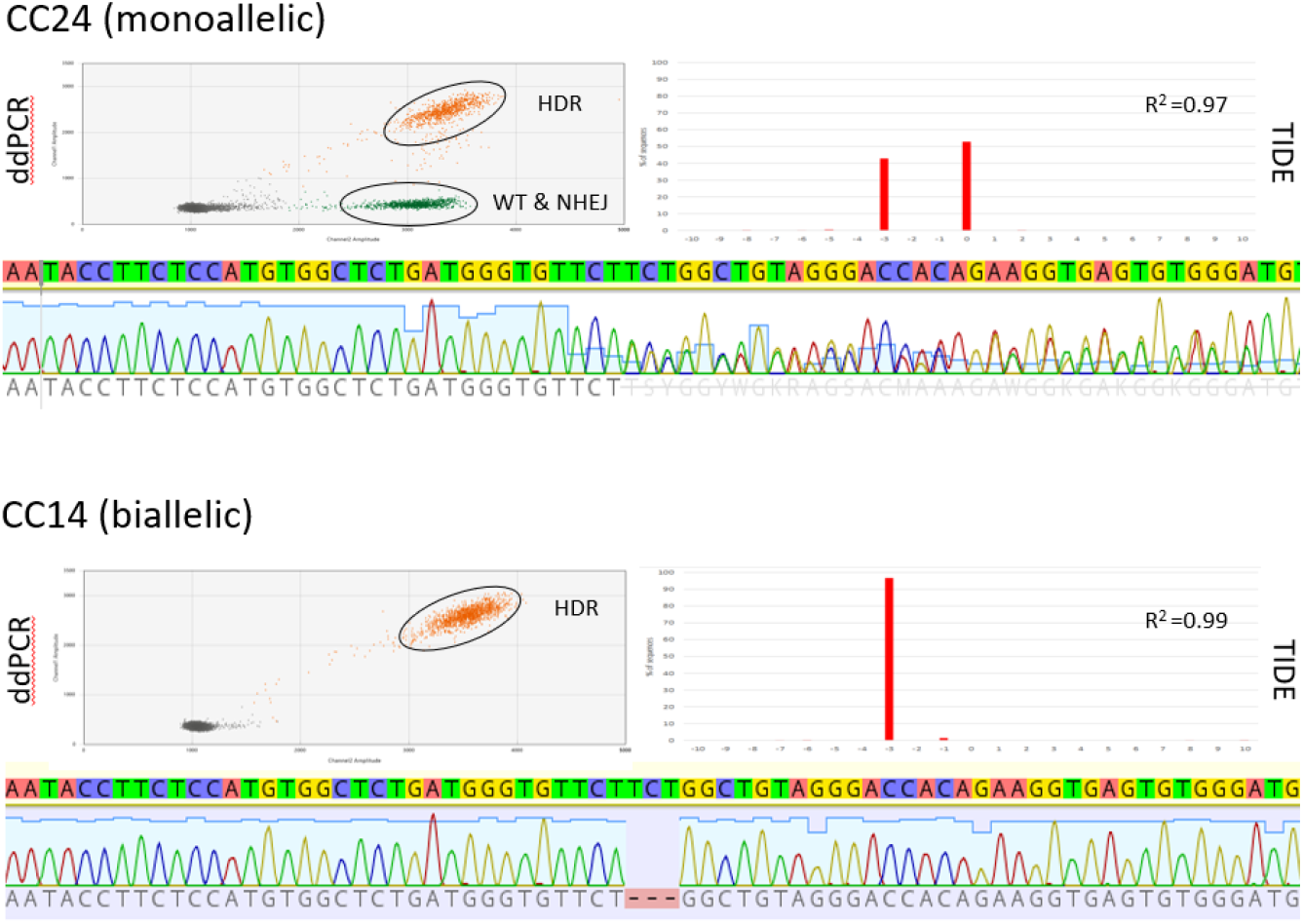
Monoallelic and biallelically edited cell clones. Examples of cell clones with a monoallelic (CC24) and biallelic (CC14) *PMEL* HDR edit. Shown are the results for their characterisation by ddPCR, sequencing and TIDE analyses. HDR: pool of droplets recognised by the HDR probe; WT & NHEJ: pool of droplets not recognized by the HDR probe indicating the presence of a WT or NHEJ allele; R^2^: coefficient of determination.

### Generation of edited calves

To determine the phenotypic impact of the PMEL mutation we used SCNT to generate calves with donor cells from the biallelic cell clone CC14 and the parental WT cell line BEF2. Following transfer of 22 (CC14) and 14 (BEF2) reconstructed embryos a total of seven pregnancies were detected at day 37 of gestation (Table 4). For each genotype, one pregnancy failed, resulting in the birth of two PMEL mutant calves and three control calves. The health of one of the PMEL calves was compromised at birth due to complication from a hydrops pregnancy. Although the second edited calf was healthy at birth, it died at the age of four weeks due to an undetected naval infection. All three control calves were also healthy at birth and developed normally.

**Table 4.** Summary of nuclear transfer results.

| Cell clone | Genotype | Embryos transferred | Pregnancies at D37 of gestation (%)<sup>a</sup> | Development to term (%)<sup>a</sup> | Development to weaning (%)<sup>a</sup> |
| --- | --- | --- | --- | --- | --- |
| CC14 | 3 bp deletion | 22 | 3 (14) | 2 (9) | 0 (0) |
| BEF2 | WT | 14 | 4 (29) | 3 (21) | 3 (21) |

<sup>a</sup> per total number of transferred embryos

### Genotype and phenotype characterisation

The genotype of the calves was confirmed by sequencing of the target region. Genomic DNA isolated from the calves revealed uniform sequences indicating the presence of two identical alleles. Both PMEL mutant calves were of the three bp deletion genotype whereas the genotype of the control calves was WT (Figure 3).

**Figure 3.**
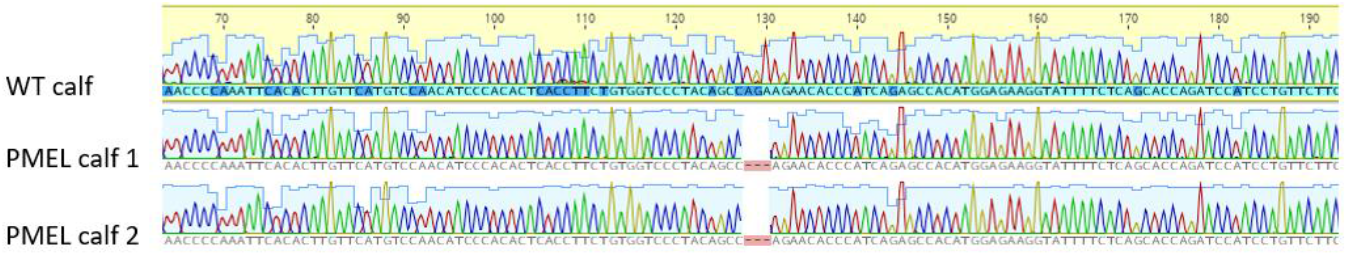
Target site sequence of the genome edited calves. Shown is an alignment of Sanger sequence results of the *PMEL* target region of one WT calf and the two mutant calves, genome edited for the p.Leu18del PMEL mutation.

Because the gRNA/Cas9 editors were delivered as plasmids, the genomic DNA of the calves was assessed for potential vector integration by PCR. PCR amplification of a 472 bp vector-specific fragment from the calves’ genomic DNA failed for both PMEL mutant calves and the three control calves and only the positive control produced an amplification product (Figure 4A). By contrast, amplification of a genomic fragment from an endogenous bovine gene was successful for all calves (Figure 4B). This result suggests that the edited calves were free of unwanted plasmid vector integrations.

**Figure 4.**
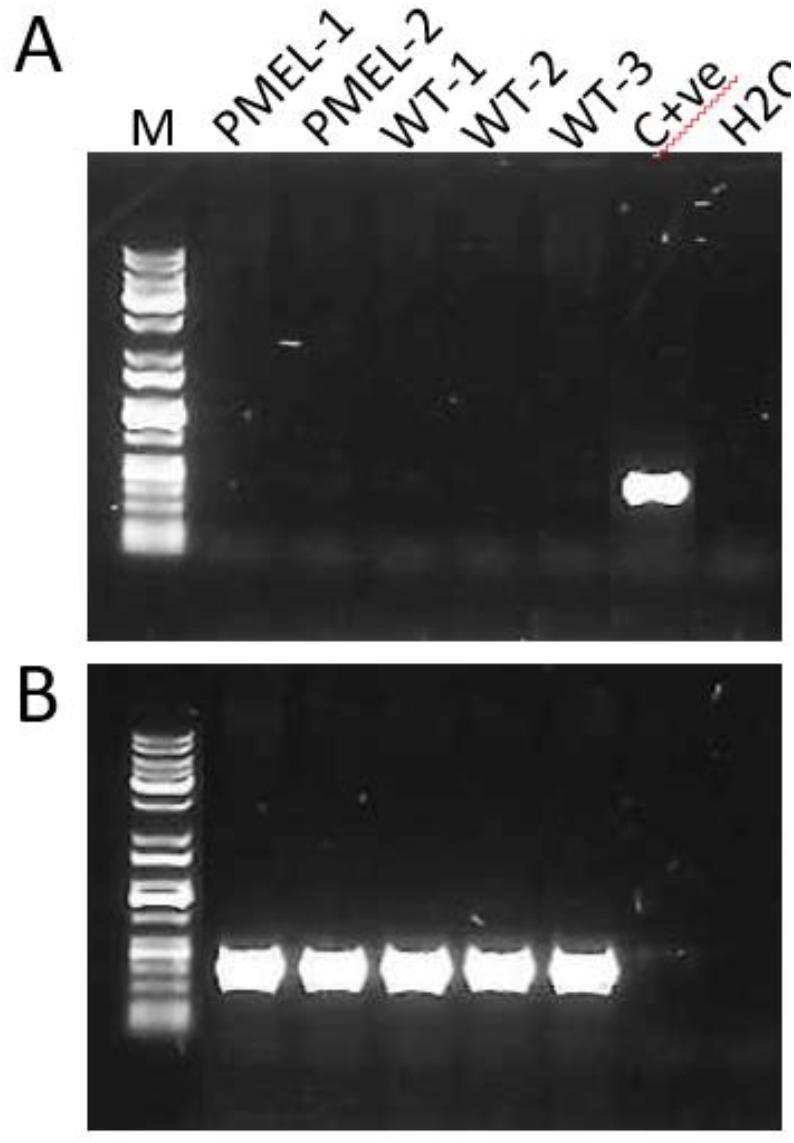
Absence of a plasmid-specific fragment in genomic DNA from edited calves. (A) Shown are amplification results for a gRNA/Cas9 plasmid-specific amplicon with genomic DNA isolated from the two edited calves (PMEL-1, 2) and the three non-edited control calves (WT-1, 2, 3). M: DNA size marker; C+ve: positive control of gRNA/Cas9 plasmid; H2O: water control. (B) The same samples analysed for the amplification of a genomic fragment specific for the endogenous *LALBA* gene encoding alpha-lactalbumin.

Both PMEL mutant calves, homozygous for the edited three bp deletion, showed a diluted coat color phenotype (Figure 5). Instead of the typical black and white coat color pattern of the Holstein Friesian cattle breed displayed by the WT control calves, the dark coat markings of the two mutant calves were no longer black but diluted into a much lighter shade of color. This resulted in a striking coat color phenotype of grey and white. Commonly, the color patterns of cloned cattle are not identical but remain very similar, which was the case for the three control calves. By contrast, the pattern of the PMEL mutant calves diverged beyond similarity of the observed pattern of the WT control clones. The markings on the face of the calves serve as an example to best illustrate this exaggerated depigmentation phenotype. Whereas the WT calves possessed a very characteristic black face with a white diamond shape on the forehead, both PMEL mutant calves had a white face. In addition, the total white areas on the coat appeared to be increased in the PMEL mutant calves compared to the WT control calves. Given that major-effect QTL for white spotting have recently been reported in NZ dairy animals (29), we characterised the status of these loci in our foundation cell lines to provide context to the apparent impacts of p.Leu18del on spotting. This analysis showed edited and control animals were homozygous for two of the three ‘white increasing’ alleles reported by Jivanji and co-workers (29), suggesting the genetic background of our animals was near ‘maximally-spotted’ as determined by these other major coat-color genes (Table S1).

**Figure 5.**
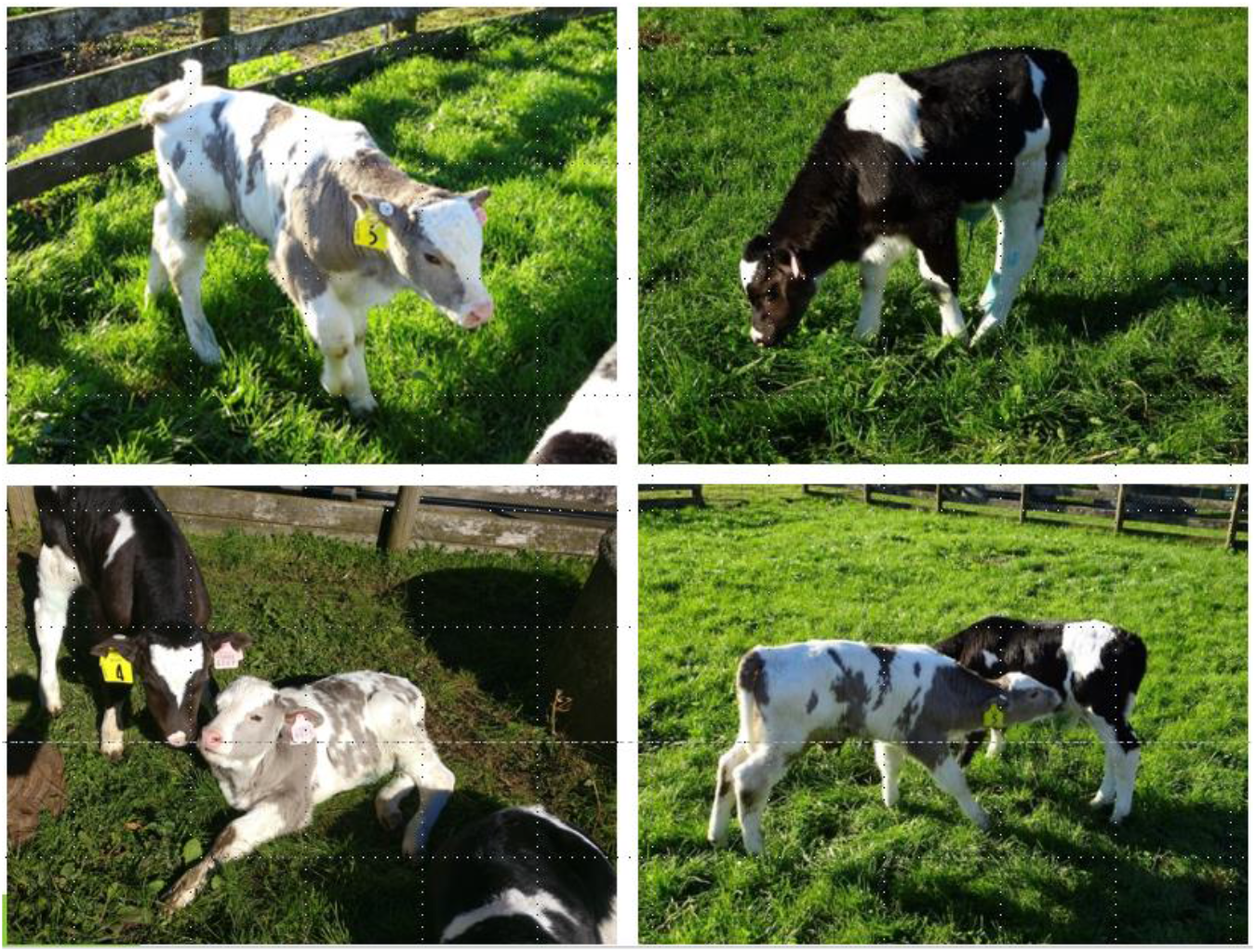
Color dilution phenotype of genome edited calf. Shown are pictures of the PMEL mutant calf with non-edited control calves for direct comparison of coat colors and distribution of white and dark markings.

To quantify the color dilution effect caused by the PMEL mutation, colorimetric measurements of hair and underlying skin color were taken from the three control calves and the viable PMEL calf. The characteristic measured was the lightness of the color expressed as L* value according to the CIELab color scheme which defines the lightness of white as 100 and black as zero. The color shade of the white hair of the PMEL calf (L* = 81.6) was indistinguishable from the white hair of the three WT control calves determined as 80.8, 83.5 and 81.3 (Figure 6). This was in stark contrast to the shade of the dark hair, with a strong increase of the lightness for the PMEL calf (L*=53.8) compared to the three WT control calves (L*=16.8, 20.5 and 16.5; P < 0.0001). To assess any associated changes in the pigmentation of the underlying skin, lightness was measured for the skin below white and dark-haired coat markings. The L* value for skin beneath white-haired markings was the same for the PMEL mutant calf (L*=72.4) and WT control calves (L*=71.7, 73.0 and 71.3). However, similar to the difference observed for the darker hair, the shade of color for the skin under the darker coat markings showed a marked difference between the mutant calf (L*=45.1) and the WT controls (L*=23.8, 33.5 and 28.3; P < 0.0001).

**Figure 6.**
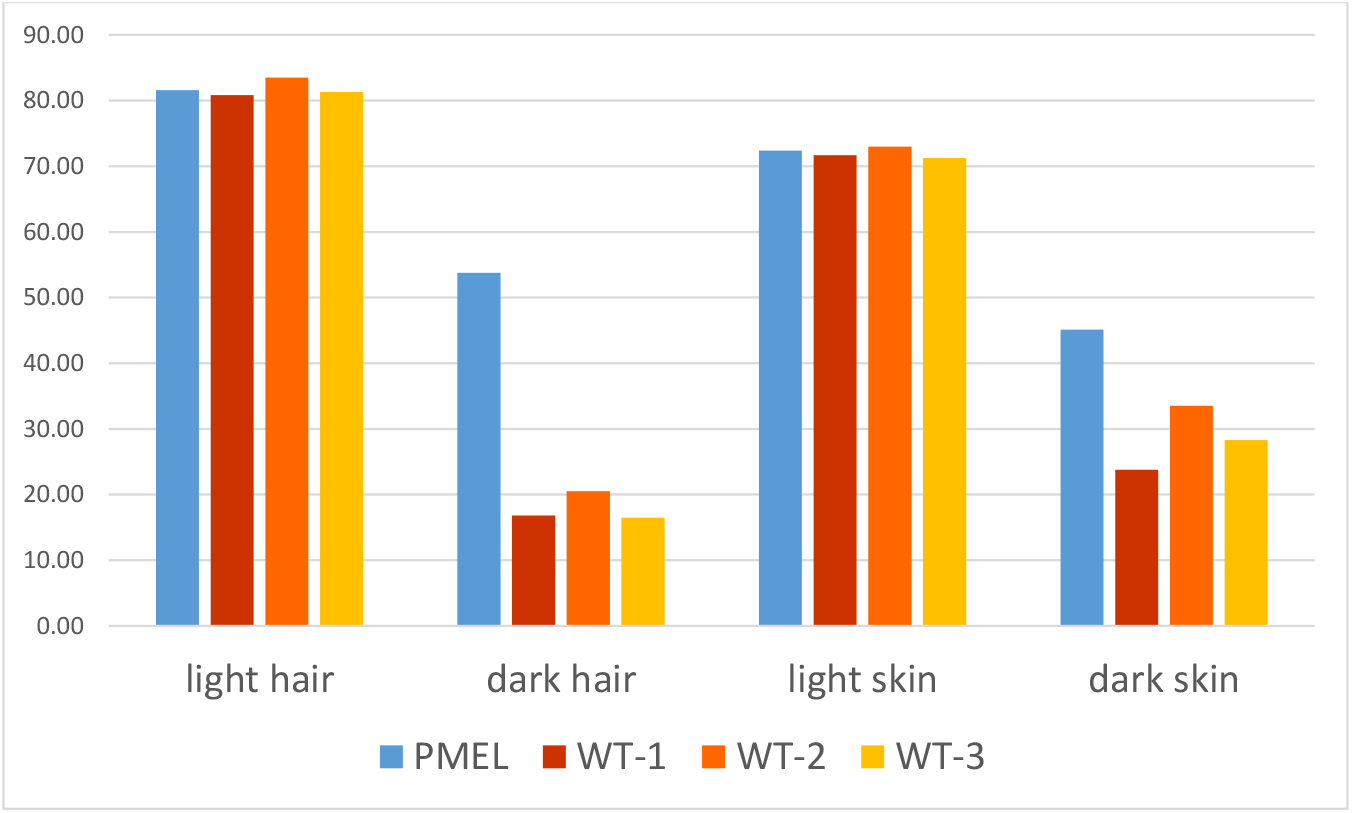
Color dilution effect on hair and skin color of edited and WT calves. Bar graphs represent the lightness of the shade of color according to the CIELab color scheme. Shown are the averages of multiple measurements at two different sites for hair and skin as indicated. PMEL: calf edited for the p.Leu18del PMEL mutation; WT: wild type control calves.

### Impact of p.Leu18del on the processing of the PMEL signal peptide

Considering the remarkable phenotypic impact of deleting a single leucine residue in the signal peptide of PMEL, we wondered about what impact p.Leu18del might have on the processing of the signal peptide, the N-terminal end and overall length of the protein.

For WT bovine PMEL (UniProt entry Q06154), the database annotation specifies a 24 amino acid (aa) signal peptide. To assess the potential impact of the deleted leucine, we applied signal peptide predictions for WT and p.Leu18del PMEL sequences (30). For both forms, presence of a signal peptide is predicted with a likelihood of 0.89 for WT and 0.85 for the mutated PMEL (Figure 7). The cleavage site for the WT is predicted between aa 26 and aa 27 (TEG-PR) which differs from the database annotation. More importantly, the cleavage site predictions for WT are also different compared to the p.Leu18del PMEL variant. For the variant, the cleavage site was predicted between aa 19 and aa 20 (VLA-VG) which also provided supporting evidence that the signal peptide of the PMEL deletion variant is processed. Hence, the deletion of leucine 18 in the signal peptide is predicted to cause the generation of an N-terminal domain that differs in length (plus six aas) from WT PMEL. These results prompted us to examine the functionality of the signal peptide in another N-terminal PMEL variant, p.Gly22Arg, that has been associated with coat color dilution in Charolais cattle (31). Similar to the p.Leu18 del mutation, the signal peptide appears to be functional with a predicted likelihood of 0.83 and cleavage site between aa 23 and 24, extending the N-terminal domain by two aas compared to WT PMEL (Figure 7).

**Figure 7.**
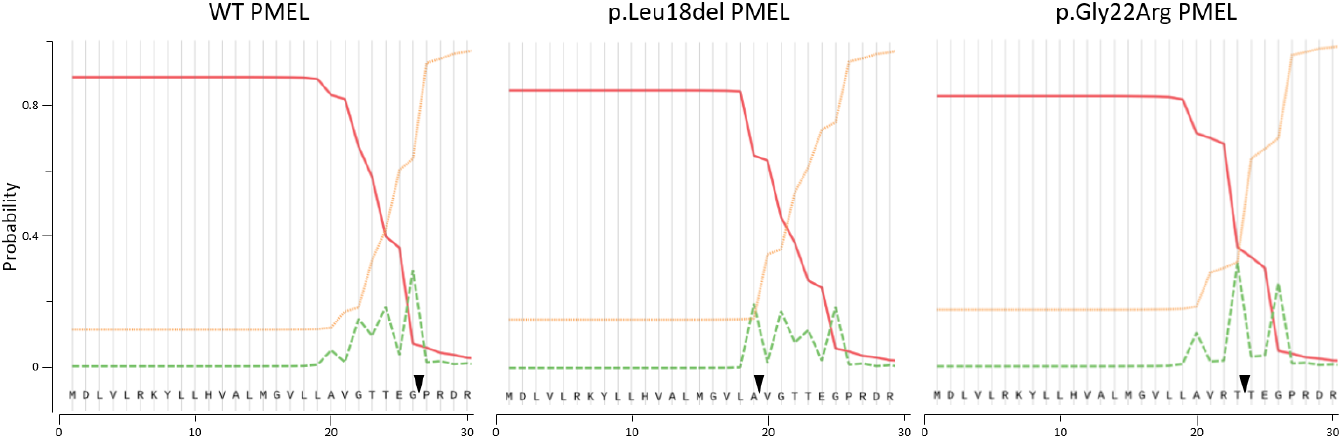
Signal peptide prediction for WT, p.Leu18del and p.Gly22Arg PMEL variants. Plotted are the probabilities reported by Signal P-5.0 for a signal peptide (solid red line), for the cleavage site (dashed green line) and for no signal peptide (dotted yellow line) in the indicated PMEL variants. Arrowheads indicated the predicted cleavage sites.

## Discussion

Initially, we designed and tested three different gRNAs for introducing the naturally occurring three bp deletion in the PMEL gene that has been associated with coat color dilution. All three showed in combination with Cas9 target site-specific cleavage activity and the ability for HDR-induced introduction of the template-specified three bp deletion following transfection into bovine cells. This also showed that HDR could be achieved without introducing an additional mutation to eliminate the PAM motif and enabled the introduction of only the naturally occurring three bp deletion. The gRNA/Cas9 editor with the highest HDR results was chosen for the isolation of edited cell clones. Although an additional PAM mutation might have increased the overall HDR efficiency, the gRNA/Cas9 editor was sufficiently discriminatory for HDR and WT-allele to allow for isolation of correctly edited cell clones.

Instead of using single cell sorting or antibiotic selection that might put cells under increased stress, we opted for manual picking of individual mitotic cells as a gentle method for the isolation of edited cell clones. A large proportion of transferred cells (75 %) successfully proliferated into expanded cell clones that were available for screening and subsequent production of edited calves. As expected from our low stringency PCR screen, rescreening of putative candidates with higher stringency identified several clones as false-positive. Even our stringent ddPCR assay using a hybridisation probe that was designed to selectively bind the correctly edited sequence showed some promiscuity. The assay also identified six cell clones, albeit at a lower (1000) than expected (2500) amplitude, that had a four bp instead of a three bp deletion at the target site (data not presented). Still, correctly edited cell clones could be readily identified by the assay and seven clones had genotypes with the precise, naturally occurring three bp deletion in the *PMEL* gene. Overall, the HDR-editing efficiency proved to be sufficiently high (7/97; 7.3 %) to isolate several cell clones that were monoallelically and biallelically edited with precision by HDR.

The edited cells were suitable donor cells for SCNT and generated two cloned calves at term, which was similar to the cloning efficiency observed with non-edited control cells. The health of one of the PMEL calves was adversely affected by the effects from a hydrops pregnancy and was euthanized shortly after birth on welfare grounds. The second calf appeared healthy and thriving initially but was lost after four weeks due to a naval infection that went undetected. Both conditions are known complications associated with the generation of cloned cattle and are assumed to be triggered by incorrect epigenetic reprogramming of the donor genome (32). Although all three control calves remained healthy, the cloning success from edited and control donor cells was not significantly different (P > 0.1). Still, edited cells were put under additional stress from single cell clonal expansion and prolonged time in in vitro culture compared to control cells which might affect their clonability. Low cloning efficiencies into viable edited calves of the cell-mediated editing approach could potentially be avoided by directly editing in vitro produced zygotes that are not compromised in their developmental potential (33). However, this approach has its own shortcomings. It lacks full control over the time and extent of editing and can generate complex mosaic genotypes, which to a certain extent can be addressed by screening embryos prior to transfer (27). Both calves had the expected edited genotype of the precise biallelic three bp deletion in the *PMEL* gene that had been confirmed for the donor cell clone used for SCNT. Because the editor was delivered by a plasmid there is a potential for unintended vector integration. Although, for a circular, supercoiled plasmid integration events are rare (34), plasmids can integrate into random genomic loci (35) or in combination with site-specific editors, such as TALENS or gRNA/Cas9 nucleases, at the target site for double strand cleavage (36, 37). The risk for such unintended integration of the molecular tools could be avoided by delivering editors as RNA molecules or as ribonucleoprotein complexes in the case of Cas9/gRNA, which can efficiently deliver editing activity into cells but are not substrates for possible integration into the genome (37). Editors have also be shown to cause potential off-target mutations due to residual binding activity and thus, cleavage activity at sites that share sequence similarity with the target site (38). However, when gRNAs are used that follow appropriate design rules, now provided by many online design tools, gRNA/Cas9 editors are unlikely to generate edited animals with significant off-target mutations (39). In a separate study, the genotype of the edited calves was analyzed in detail for any potential off-target mutation. Results confirmed that both calves were accurately edited for the intended three bp deletion and did not reveal any evidence for the presence of potential off-target mutations (40).

PMEL is a membrane protein that is exclusively expressed in pigmented cells where it is involved in the maturation of melanosomes, and pigment deposition and polymerisation in these organelles. It has a complex domain structure that has provided little insight into its function (41). Mutations in *PMEL* were associated with black pigment dilution in several species but are found in different parts of the protein. While the cattle mutations in Charolais (p.Gly22Arg) as well as Highland and Galloway (p.Leu18del) are located within the signal peptide of PMEL (18, 31), in most other species, including mouse, chicken, dog and horses, naturally occurring functional mutations are clustered in the C-terminal transmembrane and cytoplasmic domains (12–16). A more severe mutation has been generated in a chemically induced zebrafish mutant, where a premature stop codon generated a truncated protein of just over half its normal size (17). The more subtle mutations in cattle still produce a full-length protein. Assuming the predictions are correct, the mutant variants retain the functionality of the signal peptide. However, the cleavage site for the processing of the signal peptide is altered, generating a N-terminus with six and three additional amino acids for the p.Leu18del and p.Gly22Arg mutations, respectively, that are part of the cleaved off signal peptide in the WT protein. These predictions still need to be experimentally verified by determining the N-terminus of the variant forms of PMEL. If confirmed it would indicate that a seemingly minor change at the N-terminus results in at least a partial loss of functionality, revealed by the observed hypopigmentation, while the underlying molecular causes remain to be determined. Introgression of the homozygous p.Leu18del mutation into Holstein Friesian genetics resulted in a marked black pigment dilution effect. However, the dilution was not as strong compared to Highland cattle where, in homozygosity, the PMEL variant is associated with an almost white color, named silver-dun (18). This partial rescue might be explained by interactions of *PMEL* with additional genes that exist as different allelic variants in Highland and Holstein Friesian cattle. In Highland cattle, the p.Leu18 del mutation acts in a semi-dominant pattern. Whether, this holds also true for Holstein Friesian cattle need still to be addressed by determining the coat color phenotype of hemizygous animals.

In addition, there was not only a general color dilution of black markings but a marked difference in the distribution and patterning of dark and white coat markings. The PMEL mutant calf had a larger total area of white markings and a characteristic white face compared to the control calves that were predominantly black with a black face and a white diamond shape on the forehead. Given that several, major effect QTL have recently been reported for white spotting in cattle of Holstein ancestry (modulated through the *KIT, MITF*, and *PAX3* genes; (29)), these findings suggest potential epistatic interactions with these loci. Although the uniform genetic background of our study precludes a formal analysis in this regard, we genotyped tag SNPs and candidate causal mutations for these loci to provide context to the *PMEL*-derived observations, and support potential future analyses to directly test this hypothesis in alternative genetic backgrounds. This analysis showed that our study animals carried a near full complement of ‘white-increasing’ alleles, suggesting introgression on other Holstein backgrounds might yield further (though incremental) gains in depigmentation. It is noteworthy that all four of these genes interact either directly or indirectly through shared pathways controlling melanoblast migration, differentiation, proliferation and/or survival (42, 43). Epistatic interactions are therefore anticipated and might help further define the signalling relationships between these molecules and underlying melanocyte biology.

In summary, our study demonstrates the introgression of a precise *PMEL* mutation that naturally occurs in Highland cattle into both alleles of Holstein Friesian cattle. This mutation was associated with a coat color dilution phenotype in Highland cattle. When introgressed into Holstein Friesian cattle, it led to a coat color dilution effect, lightening the black coat markings to a silvery grey color. To our knowledge this is the first example that has functionally confirmed the causative status of a mutation for a coat color phenotype in cattle. The rational of lightening the coat color was to alleviate heat stress and provide adaptation to warmer summer temperatures. The edited genotype now represents a valuable model to study the impacts on heat tolerance and determine any potential effects it might have on reproductive performance, milk production characteristics and resilience to diseases associated with exposure to sunlight. Although we have demonstrated it for a dairy breed, the strategy could be readily applied to beef breeds such as Black Angus. Projected onto a global scale, even modest improvements of eco-productivity from color-diluted cattle would translate into substantial environmental benefits. Overall, our study exemplified and validated genome editing as a promising new approach for the rapid adaptation of livestock to changing environmental conditions.

## Acknowledgements

We thank Stephanie Delaney and Ruakura farm staff for dedicated animal husbandry, Fanli Meng and Pavla Turner for assistance with SCNT and Suzanne Rowe for critical reading of the manuscript. This work was funded by AgResearch and the Ministry of Business, Innovation and Employment.

