## Supplemental table 1 for "Holstein Friesian dairy cattle edited for diluted coat color as adaptation to climate change"

**Table S1**. Genotype for major-effect QTL associated with white spotting.

| QTL* | ID | Position ARS_UCD1.2 | Q allele+ | q allele# | BEF2 gen@ | CC14 gen& |
|---|---|---|---|---|---|---|
| chr2_PAX3 | rs109979909 | chr2:110817975 | A | C | AC | AC |
| chr6_KIT | rs451683615 | chr6:62557125 | A | G | AA | AA |
| chr22_MITF | rs209784468 | chr22:31651379 | A | G | AA | AA |

* QTL from Jivanji et al. 2019; positions reference tag variant or candidate mutation

+ Allele associated with increased spotting

### Allele associated with decreased spotting

@ parental cell line

& edited cell line
